# Sex enhances survival in *Paramecium*

**DOI:** 10.1101/861187

**Authors:** Amarinder Singh Thind, Valerio Vitali, Mario R. Guarracino, Francesco Catania

**Affiliations:** Institute for High Performance Computing and Networking (ICAR), National Research Council (CNR), Via Pietro Castellino 111, 80131 Naples, Italy; Institute for Evolution and Biodiversity, University of Münster, Hüfferstrasse 1, 48149 Münster, Germany

**Author notes:** These authors contributed equally (Amarinder Singh Thind, Valerio Vitali). To whom correspondence should be addressed, Francesco Catania.

## Abstract

The pervasiveness of sex despite its well-known costs is a long-standing puzzle in evolutionary biology. Current explanations for the success of sex in nature largely rely on the adaptive significance of the new or rare genotypes that sex may generate. Less explored is the possibility that sex-underlying molecular mechanisms can enhance fitness and convey benefits to the individuals that bear the immediate costs of sex. Here we show that self-fertilization can increase stress resistance in the ciliate *Paramecium tetraurelia*. This advantage is independent of new genetic variation, coupled with a reduced nutritional input, and offers fresh insights into the mechanistic origin of sex. In addition to providing evidence that the molecular underpinnings of sexual reproduction and the stress response are linked in *P. tetraurelia*, these findings supply an explanation for the persistence of self-fertilization in this ciliate.

## Introduction

One long-standing hypothesis for the evolution and maintenance of sex is that it advantageously shuffles paternal and maternal alleles thereby facilitating adaptation to changing environments **[1]** and enhancing coping with fast-evolving parasites **[2-4]**. This hypothesis fits well with the observation that a shift from asexual to sexual reproduction is initiated in many species upon exposure to environmental challenges **[5]**. However, a “*sex as diversity generator*” hypothesis might be less compelling when one considers self-fertilization. While self-fertilization can also be triggered by stressful conditions, it may give rise to offspring that are completely homozygous and genetically identical to the parent (*e.g.* in the ciliate *Paramecium* **[6]**). This implies that the selective advantage of sex may not lie only in the generation of different, potentially adaptive allelic combinations. If so, then why would selfing species—those that generate F1 copies of themselves, in particular—engage in sex upon exposure to environmental challenges? Any answer to this question needs to consider the molecular mechanisms that underlie sexual reproduction as well as existing empirical findings and theoretical arguments that sex can respond to stress **[7-9]**, is maintained upon exposure to parasites **[10-14]**, is often condition-dependent **[15]**, may be induced in response to a host immune system **[16, 17]**, may have higher fitness than asex **[18]**, and increases the rate of adaptation **[19]** upon exposure to a harsh (but not more favorable) environment **[20]**.

Manifold investigations have attempted to characterize the possible benefits of sex focusing either on how sex has become widespread despite its costs (*e.g.*, **[18, 21]**), or on the population genetic mechanisms that drive its evolution (*e.g.*, **[22, 23]**). The possibility that sex-underlying molecular mechanisms *per se* (*i.e.*, regardless of the concomitant effect of sex on genetic diversity) provide a fitness advantage remains largely neglected **[24]**. This is an important oversight given that the properties of sex reflect the interplay of intracellular and selective processes. If the capacity for sex and the organismal stress response overlap, *i.e.* have molecular components in common, then sex could be triggered when there are environmental challenges and enhance survival at the same time.

This possibility aligns with evidence that highly conserved components of the cellular stress response machinery such as heat shock proteins (HSP) play crucial roles in reproductive development and sex. Although HSP are probably best known as major players in the counteraction of proteotoxic damage, these proteins can play other biological functions e.g. they regulate mitotic cell division **[25]** and stimulate the immune response **[26, 27]**. HSP genes are also expressed physiologically during oogenesis and/or spermatogenesis across a wide range of species **[28-37]**. Members of the HSP70 family are required for the process of meiosis: HSP70 is key to the formation of the synaptonemal complex in vertebrates **[38-40]** and in protozoans **[41]**. Furthermore, HSP70 is required for fertilization in vertebrates **[42]** and protozoans **[43]**. Because of the strong molecular ties that link HSP to sexual reproduction, if sex-related intracellular processes can truly enhance fitness in the context of environmental alterations, then HSP with reproductive functions may contribute to the advantage of sex.

But what might this advantage be? For one, HSP with reproductive functions (HSP_R_) might help activate HSP with stress-related functions (HSP_S_) upon exposure to environmental stress, benefitting either the individuals that undergo sexual reproduction, or their resulting offspring, or both. This hypothesis is consistent with observations in the ciliate *Tetrahymena thermophila*, where the cytosolic HSP70 SSA5 chaperone has a crucial role in pronuclear fusion during fertilization and is also expressed in response to heat stress—without directly contributing to cell survival however **[43, 44]**. In a non-mutually exclusive fashion, the expression of HSP_R_ might also enhance survival by reducing sensitivity to environmental alterations. These circumstances are reminiscent of the diapause: a widespread and reversible state of dormancy that is achieved through the involvement of HSP and insulin/insulin-like signaling **[45-51]**. Finally, HSP_R_ might facilitate sexual reproduction in stressful environments. Elevated levels of environmental stress in early life might prematurely activate HSP_R_ (letting alone HSP_S_), and in so doing ultimately accelerate reproductive development. The resulting association between early life stressors and accelerated sexual maturation is consistent with a wealth of observations **[52-54]** and involves, once again, the modulation of insulin/insulin-like signaling **[55, 56]**. It is worth emphasizing that stress over an organismal lifetime can be experienced even without ecological changes. Mechanisms and molecular products that are commonly associated with a stress response and induce expression and accumulation of HSP70 (*e.g.*, apoptosis, reactive oxygen species) have been linked to development and sexual reproduction **[57-62]**.

*Paramecium* is an ideal organism to evaluate the significance of the foregoing hypothetical scenarios. One of the best studied species of this genus, *P. tetraurelia*, can reproduce asexually, via binary fission, or sexually via meiosis followed by either cross-fertilization (conjugation) or self-fertilization (autogamy) **[6]**. Autogamy in *P. tetraurelia* occurs in unpaired cells, is induced by starvation, and generates fully homozygous individuals **[63-65]**. More specifically, *P. tetraurelia* contains two types of nuclei, a micronucleus (with germinal functions) and the macronucleus (with somatic functions). When exposed to a sufficiently long period of food restriction, the macronucleus of *P. tetraurelia* undergoes fragmentation and is gradually reabsorbed via apoptosis- and autophagy-like processes. The two remaining micronuclei undergo two meiotic divisions producing eight haploid nuclei, one of which is preserved and divides mitotically to form genetically identical gamete nuclei that unite to form a completely homozygous fertilization nucleus. This latter nucleus then devides twice, producing new germline diploid and somatic polyploid nuclei **[6]**. Autogamy rejuvenates *P. tetraurelia* **[65]**. Furthermore, *P. tetraurelia* is more likely to undergo autogamy when sufficiently old: at comparable levels of starvation, ∼27 fission-old *P. tetraurelia* is far more likely to undergo autogamy compared to cells that are ∼15 fissions away from the last fertilization **[66]**. Interestingly, the level of starvation required for initiation of autogamy cells scales negatively with clonal age **[67]**. Besides, the length of autogamy immaturity is linked to clonal life span and rate of cell division **[68-71]**. The reasons for these relationships remain largely unknown in *P. tetraurelia*. In multicellular systems the insulin/IGF-1 signaling pathway plays a key role in shaping similar relationships **[72]**.

Here, we leverage *P. tetraurelia* to ask whether self-fertilization can enhance survival probability in the absence of new genetic variation. We show that fully homozygous self-fertilizing *P. tetraurelia* cells display an increased heat shock resistance, and so do also cells that have recently achieved competence for sex. A time-course transcriptomics study alongside additional phenotypic essays provide first insights into the molecular mechanisms that underlie this survival advantage. We find (i) that the observed enhanced stress resistance is negatively coupled with cell proliferation—a relationship that may be partly shaped by the withdrawal of growth factor(s)—and (ii) that self-fertilization, onset of sexual maturation, stress response, and cell proliferation are interlinked in *P. tetraurelia*. These findings suggest that sex in *Paramecium* may be advantageous even in the absence of new genetic variation, and that HSP-related molecular mechanisms are likely contribute to the preservation of self-fertilization in *P. tetraurelia*.

## Results

### Sex yields a survival advantage in the absence of new genetic variation

We reared three genetically uniform mass cultures of *P. tetraurelia* from a self-fertilization state (day 0) up to ∼33 fission-old cells (day 8) in parallel. The vegetative mass cultures were fed constantly and maintained at a density of ≤750 cell/ml to minimize stress. In these experimental conditions, the rate of cell proliferation between days (other than day 0) varies to some considerable, but not statistically significant, extent (Kruskal-Wallis test, *P =* 0.10), and amounts to an average of ∼4 fissions/day (**Figure 1A**). We used these clonally aging mass cultures to investigate the association between autogamy competence and stress resistance.

**Figure 1.**
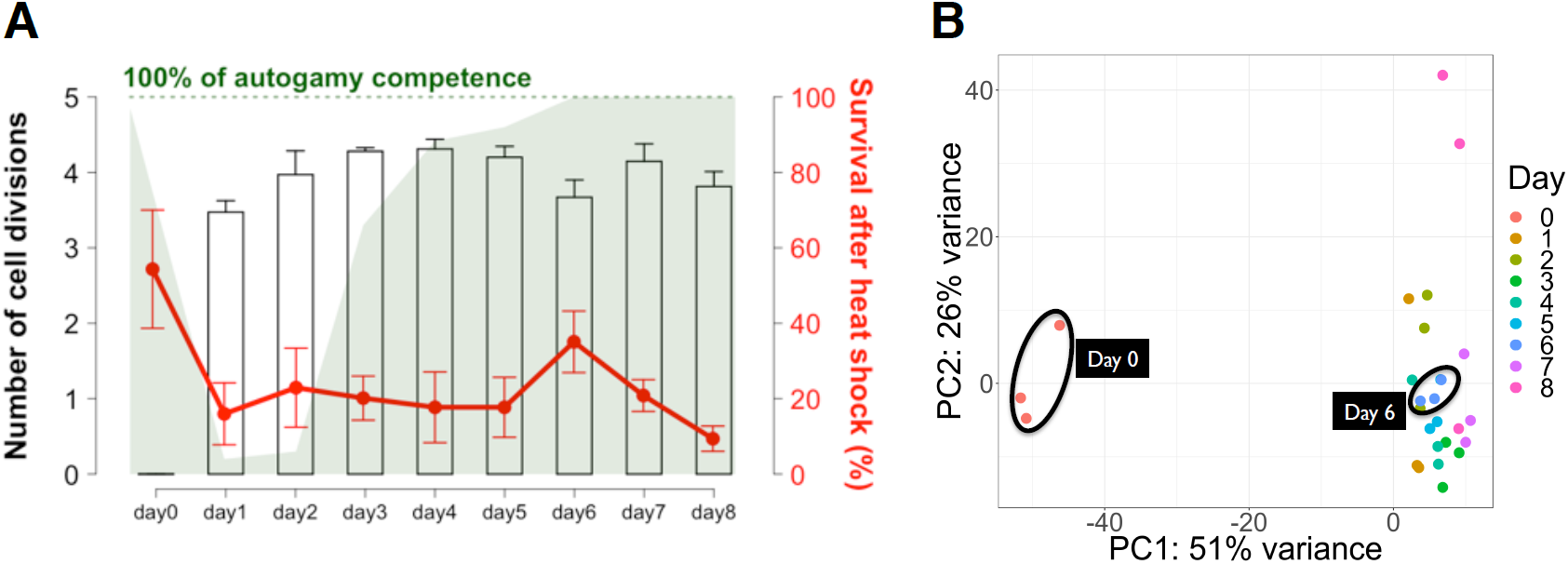
Phenotypic and transcriptomic profiles of *P. tetraurelia* cell cultures during nine consecutive days post and including self-fertilization. (**A**) Bar plot shows the number of cell divisions *per* day (average and standard errors; three replicates). The *per*-day degree of autogamy competence is reflected by the height of the green area. Line plot shows cell survival (*i.e.*, active swimming) after a 43°C heat shock (90 seconds) and 24-hour recovery time in a nutrient-rich medium at 25°C (average and standard errors). (**B**) Principal component analysis of the *P. tetraurelia* transcriptome (nine successive windows of time × three replicates).

Every day, we screened each of the mass cultures for autogamy competence. We collected aliquots of >50 cells *per* cell culture, on average, subjected them to three consecutive days of starvation, and discarded them once the phenotype had been recorded. The vast majority of day0 cells show signs of autogamy (97% ± 1.73%; mean and standard error, respectively). As expected, the fraction of cells that are capable of undergoing autogamy increases with clonal age (**Figure 1A**). There is a dramatic rise in the percentage of autogamy-competent cells from day 2 to day 3 (6 ± 1 *vs* 66 ± 4.6), which is followed by a steady upward trend (day 4: 88.3 ± 2.6; day 5: 92 ± 2.1). At day 6, all the cell cultures display 100% autogamy competence, for the first time after the last fertilization. This level of autogamy competence holds for the well-fed day7 and day8 cells. It is worth noting that the onset of full sexual competence could be immediately followed by self-fertilization in nature when nutrients are scarce.

Additional aliquots of 96 cells were collected daily from each of the mass cultures for a stress test. These cells were exposed to a heat shock of 43°C for 1.5 min, and then stored separately for 24h in a nutrient-rich medium before being screened for survivors and ultimately discarded. We detected a considerable level of variation among *P. tetraurelia* cells collected over the nine experimental time intervals. More than 50% of day0 cells (54.3% ± 15.7) survived the thermal shock, >2.5 times more than the average fraction of survivors recorded for the vegetative samples (∼20% ± 13.0; Wilcoxon Rank Sum test, *P*<0.05) (**Figure 1A**). Among the cells in vegetative state, day6 cells display a percentage of survivors (35.1 ± 8.2) that is 1.5 times lower compared to day0 cells, but ∼2 times higher than the counterpart calculated for the remaining vegetative samples (17.8 ± 11.6) (Wilcoxon Rank Sum test, *P*<0.05) (**Figure 1A**). The enhanced resistance to heat displayed by day6 cells is associated neither with genetic variation nor with clonal age, as evidenced by the unremarkable survival rates of day7 and day8 cells. It is instead associated with the onset of full competence for sex; at day 6, all the cell cultures display 100% autogamy competence for the first time after the last fertilization (**Figure 1A**).

### Transcriptional changes during sexual development are coupled with increasing levels of endogenous stress

To gain insight into the association between heat resistance and autogamy competence, we investigated the transcriptomes of the twenty-seven mass cultures (9 × 3 replicates) that were screened for these traits. A PCA reveals differences in the transcript abundance of self-fertilizing and vegetative cell cultures, as expected (**Figure 1B**). We also detect a tendency for day0 and day6 cell cultures to contain more genes with comparable levels of expression relative to other pairwise combinations (**Figure S1**). Of 32,252 genes with non-zero expression values across each of the experimental days, 1037 and 1145 have increasing and decreasing levels of expression, respectively, over the eight vegetative time points (Pearson’s *r, P*<0.05). These genes are candidates for regulating sexual maturation and reproductive processes as well as lifespan in *P. tetraurelia* (**Table S1**). “*Cell redox homeostasis*” is among the most enriched biological processes for the genes with steadily increasing expression levels (**Table S1, Figure S2**). Among these genes, Hsp70Pt04 has alongside Hsp70Pt01 and Hsp70Pt03 the highest level of sequence similarity to the ciliate *Tetrahymena thermophila*’s HSP70 SSA5 (**Figure S3**). SSA5 is a key protein for pronuclear fusion during fertilization and is also expressed during development and in response to heat-stress **[43]**. Similarly, SSA5 homologs in *P. tetraurelia* are expressed throughout development (**Figure S3**) and are inducible after heat shock exposure (Hsp70Pt03 and Hsp70Pt04; **[73]**) or up-regulated after a 3-week long period of starvation (Hsp70Pt01; **[74]**). Hsp70Pt01 also displays elevated expression levels at meiosis **[75]**, together with seven members of the HSP70 family (**Figure 2**). These observations raise the possibility that in *P. tetraurelia*, like in *Tetrahymena*, HSP70s may also play a role in sexual reproduction and respond to environmental stress at the same time.

**Figure 2.**
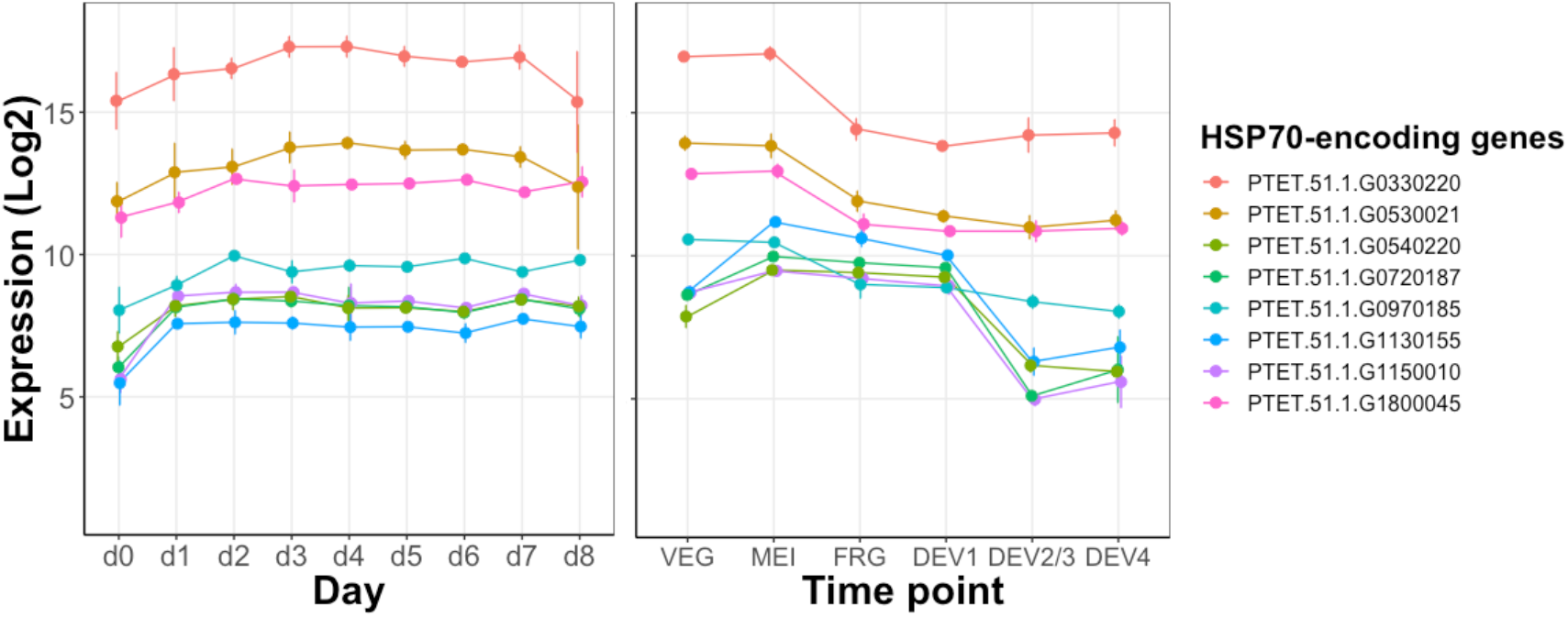
HSP70 genes expressed during vegetative growth (left, this study) and with comparable or elevated expression levels during self-fertilization (right, [75]); *e.g.* PTET.51.1.G0330220’s expression is comparable between the vegetative (VEG) and the meiotic (MEI) stage and drops before (DEV1), the earliest stage at which a significant proportion of cells has visible macronuclear anlage.

### A trade-off between stress resistance and cell division

When inspecting genes that are differentially expressed at consecutive time points or with extreme transcription activity at a given time point (**Table S1**), we uncover significantly down-regulated biosynthetic activities in day 0 and day 6 (**Figure 3, Figures S4** and **S5, Tables S2** and **S3**). This may be unsurprising for day0 cells, which neither feed nor divide. However, it is unexpected for day6 cells. Not only are these cells cultivated in a nutrient-rich environment, they also divide albeit at a marginally reduced rate compared to day5 cells (3.7 ± 0.23 *vs* 4.2 ± 0.15 divisions; paired t-test, *P*<0.05) (**Figure 1A**). When compared to day5 cells, day6 cells up-regulate a handful of biological processes such as a response to oxidative stress, whereas they down-regulate genes involved in endocytosis and components of the phagosome, among others. Additionally, our data point to a regulated withdrawal of one or more growth factors in day6 cells (**Figure S6**). These observations indicate that nutrient uptake may be hindered in day6 cells despite the nutrient-rich conditions—a state that we dub pseudo-starvation. Changes in response to oxidative stress and expression of phagosome components are also observed in the fast proliferating day3 cells (4.3 ± 0.05 divisions), however in the opposite direction. Overall, these findings suggest that levels of mitotic cell division and stress-response may trade-off in *P. tetraurelia*.

**Figure 3.**
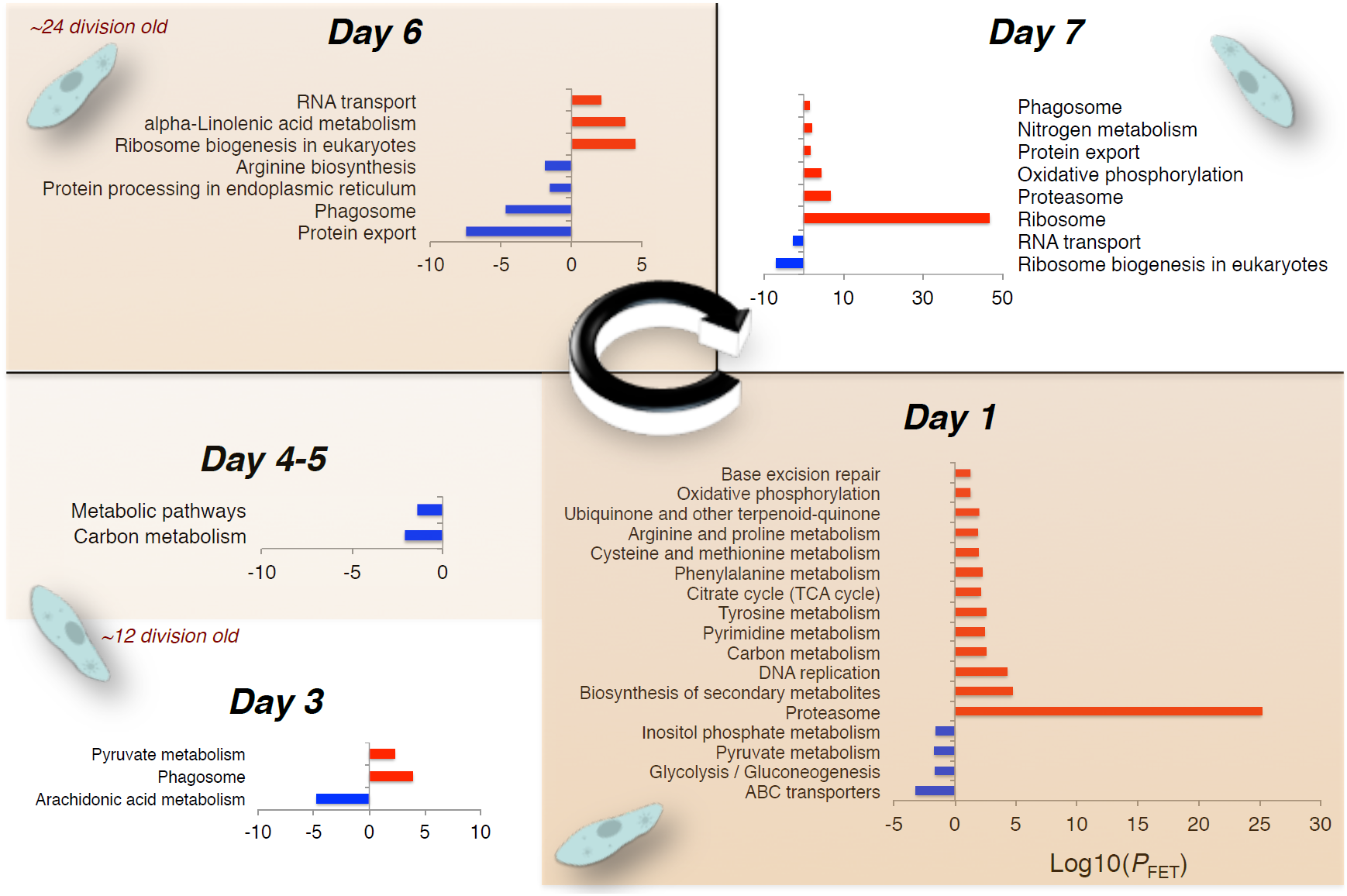
Enriched molecular pathways during the first ∼33 cell divisions post-self-fertilization. Reported enrichment (Fisher’s exact test, *P<*0.05) is relative to the previous day (e.g. Day 1 relative to Day 0) and is estimated via the functional annotation tool DAVID **[81]** and the Kyoto Encyclopedia of Genes and Genomes **[82]**.

We also found that day1 cells significantly up-regulate an excess of HSP70 chaperones (13 observed *vs* ∼4 expected; *Z* = 3.48, *P*<0.001; **Table S1**). Given that day1 cells resume proliferation after the non-dividing stage of self-fertilization, this observation aligns with the view of HSP70 chaperones as active regulators of mitotic cell division **[25]**. Two of the HSP70-encoding genes that are up-regulated in day 1 are also up-regulated in day7 cells. These two genes (PTET.51.1.G0720187, PTET.51.1.G1150010), alongside their two ohnologs, are the only HSP70s to show similarly elevated expression levels both during and between meiosis and fertilization in *P. tetraurelia* (**Figure 2**), indicating that these HSP70s might play a role in mitosis as well as meiosis, in line with previous reports **[76** and references therein].

Lastly, although day0 and day6 cells show an elevated heat shock resistance, we detected no enrichment of up-regulated stress response genes in these cells. Even more surprisingly, HSP70-encoding genes are significantly down-regulated (rather than up-regulated) in day0 cells more often than in day1 cells (13 down-regulated *vs* 1 up-regulated; **Table S1**). We interpret these observations as indicating that HSP70s already occur in day0 (and day6) cells. Following a developmentally regulated decrease of nutrient uptake, HSP that are normally involved in mitosis **[25]** might be redeployed as cytoprotectants. Consistent with this hypothetical scenario, HSP can regulate negatively the transcription of their corresponding genes **[77-79]**. Additionally, the inhibition of cell cycle genes leads to tolerance towards environmental stress in the nematode *C. elegans* **[80]**.

### Model: sex competence, mitotic progression, and stress resistance are mechanistically linked in *P. tetraurelia*

The preceding time-course study indicates that HSP with reproductive functions (HSP_R_) accumulate during vegetative growth and might respond to environmental stress in concert with HSP with stress-related functions (HSP_S_). Additionally, HSP that engage in the regulation of mitotic cell division might be redeployed as cytoprotectants when cell division is down-regulated. Taken together, these indications generate a prediction that recapitulates the observations depicted in **Figure 1**, *i.e.*, HSP_R_ and HSP_S_ shoud be more likely to co-occur in cells that divide slowly (day6) or not at all (day0). In other words, reduced levels of cell proliferation—which are controlled by nutritional input—should contemporaneously hasten the acquisition of sexual maturity and enhance stress resistance. We tested these predictions.

We examined the rates of sexual maturation in *Paramecium* cells whose proliferation is sustained by two prey bacterial species, *Enterobacter aerogenes* (EA) and *Alcaligenes faecalis* (KL), which promote different growth rates (Wilcoxon rank-sum test, *P*<0.001) (**Figure 4A**, top panel). Consistent with our prediction, we detected a significant positive correlation between cell proliferation and the duration of autogamy immaturity (*ρ* = 0.71, *P*<0.001), with the slow-growing KL-fed *P. tetraurelia* cells reaching sexual maturity earlier than the fast-growing EA-fed cells (Wilcoxon rank-sum test, *P*<0.01) (**Figure 4A**, right panel).

**Figure 4.**
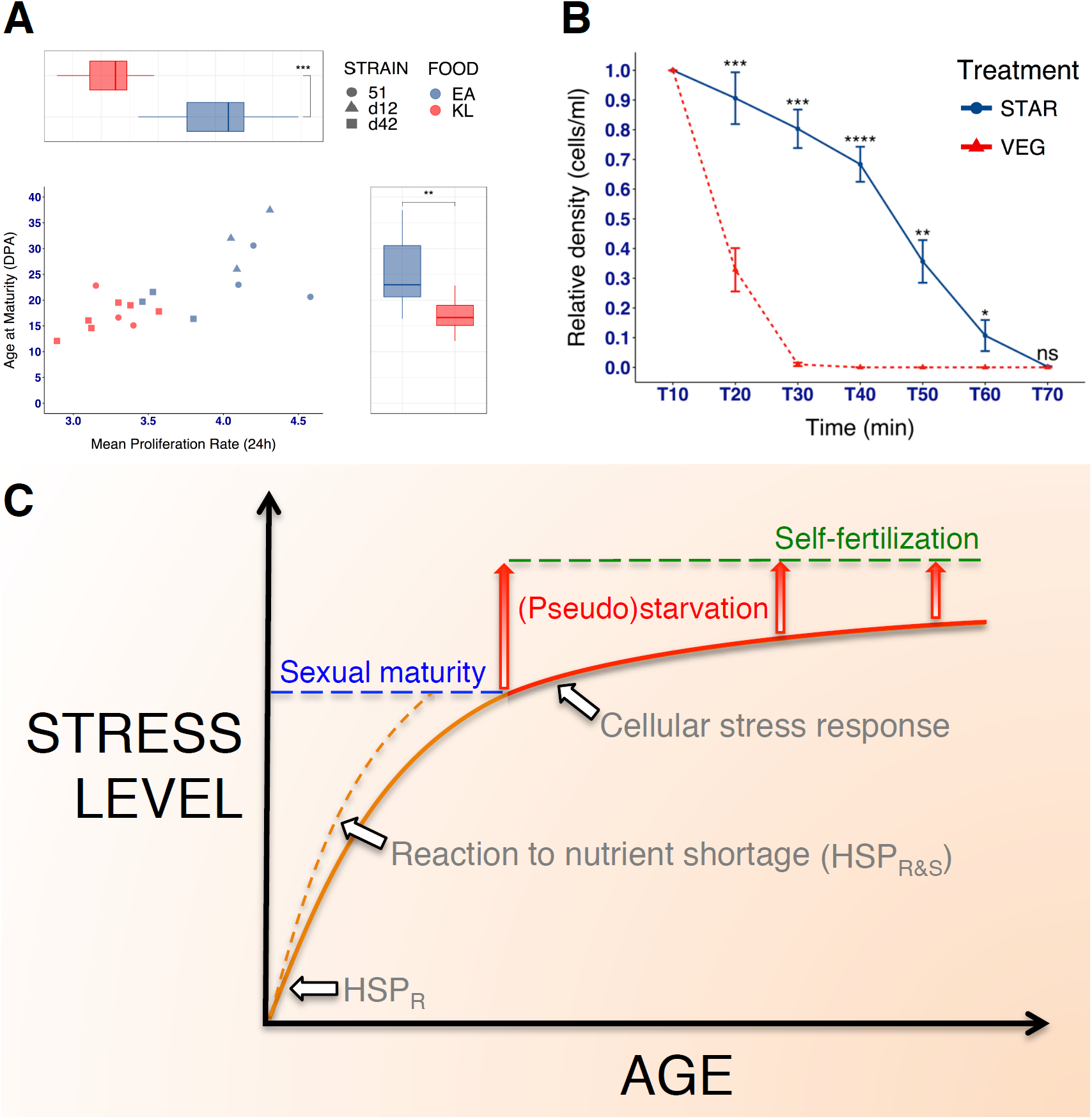
Stress response, sexual maturation and reproduction, cell proliferation, and nutritional input are mechanistically linked. **(A)** Low-quality food accelerates sexual maturation. The box plot at the top compares the main proliferation rate of cells fed with EA and KL. The box plot to the right compares the age at maturity between the two food-quality groups. DPA, Divisions Post-Autogamy. **(B)** Dietary restricted cells resist heat shock. Nutritionally deprived cells show prolonged survival during uninterrupted exposure to 40°C, whereas actively proliferating cells expire quickly. Plot shows the mean culture density +- the standard error of the mean. Mean differences were tested with a Wilcoxon signed-rank test. ns: P > 0.05; *: P ≤ 0.05; **: P ≤ 0.01; ***: P ≤ 0.001; ****: P <= 0.0001. **(C)** Model for the effect of nutritional deprivation on sexual maturation, stress response, and self-fertilization in *P. tetraurelia*. HSP with reproductive functions (HSP_R_) accumulate physiologically during development (solid orange curve) and favor sexual maturation. Nutrient shortage in the early post-autogamy period may enhance the expression of HSP_R_ and HSP with stress-related functions HSP_S_ (dashed orange curve), thereby reducing the HSP_R_ shortfall in sexually immature *P. tetraurelia* cells. This is predicted to concurrently increase the ability to cope with stress and to speed up sexual maturation (lowering the age of sexual maturity). Nutrient shortage is also required to initiate self-fertilization. A state of pseudo-starvation in sexually mature cells (arrows with red gradient, lower portion) is a first (reversible) step in the process of self-fertilization. This cellular state of regulated reduced nutrient uptake is coupled with (and possibly prompted by) a stress reaction to cell-intrinsic events. Consequently, pseudo-starving cells are particularly resistant to environmental challenges. In nutrient-limiting conditions, the pseudo-starvation state can transition into a state of full starvation, with a consequent intensification of cellular stress reactions (arrows with red gradient, upper portion). In contrast, the pseudo-starvation state is rapidly lost in nutrient-rich conditions, where normal feeding resumes. It is predicted that cells undergo a new state of pseudo-starvation after some unspecified number of divisions. As cells grow older, the cellular stress response increases and the level of starvation that is required for initiating self-fertilization decreases. It is predicted that in sufficiently (clonally) old *P. tetraurelia* cells self-fertilization is possible even in non nutrient-limited conditions.

Next, we tested whether reduced levels of cell proliferation enhance stress resistance. We compared the response of young, dietary restricted, cells and exponentially growing cells of comparable age to prolonged heat shock (**Figure 4B**). Studying young *Paramecium* cells with no or little competence for self-fertilization helps decouple the effects of dietary restriction from those of the sexual process. We detect significant pro-survival effects of nutritional deprivation (**Figure 4B**). During the heath shock bout, dietary restricted cells survived on average twice as long (long rank test, *P*<0.001), with ∼10% of the culture still alive beyond 1h of exposure (exponentially growing cells did not survive past 30 min).

Overall, these observations are encouraging. They are compatible with a model where stress resistance, autogamy maturation, and cell proliferation are mechanistically linked in *P. tetraurelia* and call for additional investigations.

## Discussion

The persistence and pervasiveness of sexual reproduction, despite its substantial time and resources requirements, continue to dumbfound biologists. The classical explanation for the success of sex is that it creates genetic variation that enhances the offspring’s chances of survival in stressful environments, or that it purges deleterious recessive alleles.

Here, we capitalize on knowledge offered by previous molecular and cell biology studies and leverage the propeties of the ciliate *Paramecium* to provide integrative answers to three questions. How can sex—self-fertilization, in particular—persist in eukaryotic species where it generates no genetic variation? Why does stress induce sex? And last, why do individuals bear the costs of sex even when they do not immediately or directly benefit from its putative benefits? Our observations suggest that the molecular mechanisms that underlie sex and stress response overlap in *Paramecium*. As a consequence, not only can stress facilitate sex in this ciliate, but sex can also take place in a molecular environment that promotes stress-resistance. Based on our findings, we propose that the coupling between stress-related and sex molecular machineries contributes to the persistence of self-fertilization in *P. tetraurelia*, a ciliate whose level of intraspecific genetic diversity **[83]** and *per* generation base-substitution mutation rate **[84]** are exceedingly low.

Our observations further suggest that stress response, the onset of sexual competence, cell proliferation, and nutritional input are mechanistically linked in *P. tetraurelia* (**Figure 4C**). We propose a model where a reduced nutrient uptake may dually speed up the acquisition of the capacity for sex and increase the ability to cope with stressors. When sufficiently prolonged, nutrient shortage can favor self-fertilization while protecting the cell against environmental stressors. These dynamics raise the possibility that even young, sexually immature, *P. tetraurelia* cells may develop the capacity for sex when they are exposed to a sufficiently prolonged nutrient shortage. This expectation is consistent with previous observations **[67]**.

The degree to which the sex-coupled survival advantage observed for *Paramecium* extends to other organisms warrants further investigation. Much can also be gained from examining the discrepancies that might emerge between *Paramecium* and other species. For example, the heat shock response (HSR) collapses with the onset of reproductive maturity in *C. elegans*, leaving adult individuals more vulnerable to stress **[85]**. This HSR collapse is maintained when the entire gonad is absent, whereas it disappears, and in fact the HSR is enhanced at the organism level, when germline stem cells (GSCs) are removed **[85]**. This suggests that *C. elegans*’ somatic gonadal cells, which play important roles in GSCs maintanance as well as in germ cell proliferation and differentiation **[86]**, produce pro-HSR signals that counterbalance the GSCs signals **[87, 88]**. While one has to draw cautious parallels, the somatic nucleus of *Paramecium* could be viewed as the somatic gonadal cells in *C. elegans*. In this light, the enhanced stress resistance that we observe for day0 and day6 cells could reflect a transiently unbalanced signalling that favors the somatic nucleus. Based on our findings, *Paramecium* could achieve this unbalanced signalling by down-regulating the nutrient-sense signaling and thus cell growth. Similar mechanisms could shape similar responses in *C. elegans* **[80, 89]**. At any rate, because of the deep evolutionary conservation of the stress response and sex molecular machineries, our study urges caution over swiftly invoking genetic diversity to explain increased stress tolerance in sexually produced offspring. Indeed, because HSP70-mediated stress tolerance may be passed on to successive sexual generations even in the absence of the parental stressor **[90]**, enhanced stress tolerance in sexually produced offspring might reflect an environmentally-induced phenotype that is trans-generationally inherited **[91]**, rather than a genetically controlled trait.

Last, the model that we propose for the maintenance of sex in *P. tetraurelia* might help shed light on the evolutionary origin(s) of sex in ciliates and perhaps other eukaryotes. Given that heat shock factors have established roles in meiosis and fertilization and that the stress response is ancient and ubiquitous—HSP expression is a universal response to a variety of stresses (*e.g.*, physical, physiological)—whereas sex is not necessary for reproduction, it is tempting to speculate that the sex machinery might have evolved from, and be partly modulated by, the heat-shock signaling pathway or, more in general, the integrated stress response **[92]**. If so, the evolutionary success of sex could partly rely on the strong and pervasive selective forces that preserve the cellular defense mechanisms.

In conclusion, we update current theories on sex benefits by illustrating the possibility that sex can confer a survival advantage in the absence of new genetic variation. Our observations in *Paramecium* extend the dominant paradigm. They suggest the existence of molecular connections between sex and stress response through which sex-performing unicellular organisms may gain direct fitness benefits.

## Methods

### Time-course experiment

#### Paramecium strain and culture conditions

Three independent polycaryonidal mass cultures of the *Paramecium tetraurelia* stock d12 **[93]** were used in this experiment. These cultures were made of fully homozygous cells, which grow on Cerophyl Medium (CM; Taylor and Van Wagtendonk 1941, 0,25% w/v Wheat Grass Powder, GSE-Vertrieb GmbH) bacterized with *Enterobacter aerogenes* Hormaeche and Edwards (ATCC® 35028TM). For bacterization, CM was inoculated (1:100 v/v) with a dense suspension of *E. aerogenes* (OD_600_=0.5) and incubated at 30°C over night under gentle shaking (120 rpm). Culture medium was prepared every 48h and supplemented with 0.5µg/ml stigmasterol (Art.No. 9468.1, Carl Roth GmbH) before use. To ensure monoxenic conditions, cells that were used for the experiment were taken from a stock, transferred into depression slides, and washed nine times in mineral water (Volvic®) before the beginning of the time-course study.

We produced autogamous mass cultures as starting material for the time-course experiment (Day 0, **Figure 5**). Briefly, stocks cells of unknown age were taken through additional ∼30 divisions to maximize autogamy reactivity and allowed to starve naturally at 25°C for three days to stimulate mass autogamy. Following a first expansion phase and unless otherwise indicated, log-phase cells were mass cultured daily in 250 ml of bacterized CM at the standard cultivation temperature of 25°C. Aliquotes of cells (30 cells/ml) from the previous day were used daily to seed the progressively (clonally) older mass cultures. Cell density never allowed to exceed 750 cells/ml. For each of the 27 mass cultures (9 days × 3 replicates), aliquotes of progressively older cells were obtained daily to perform a heat shock tolerance test (see below) and an autogamy reactivity test (see below) between 0 and ∼30 Divisions Post Autogamy (DPA) (**Figure 5**). Cell age expressed in DPA was calculated as the cumulative daily growth rate up to day *n* according to the following equation:

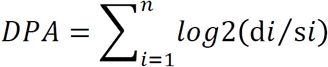

where d is the final cell density, s is the seeding cell density and log2(d*i*/s*i*) is the growth rate for day *i* expressed in divisions per day.

**Figure 5.**
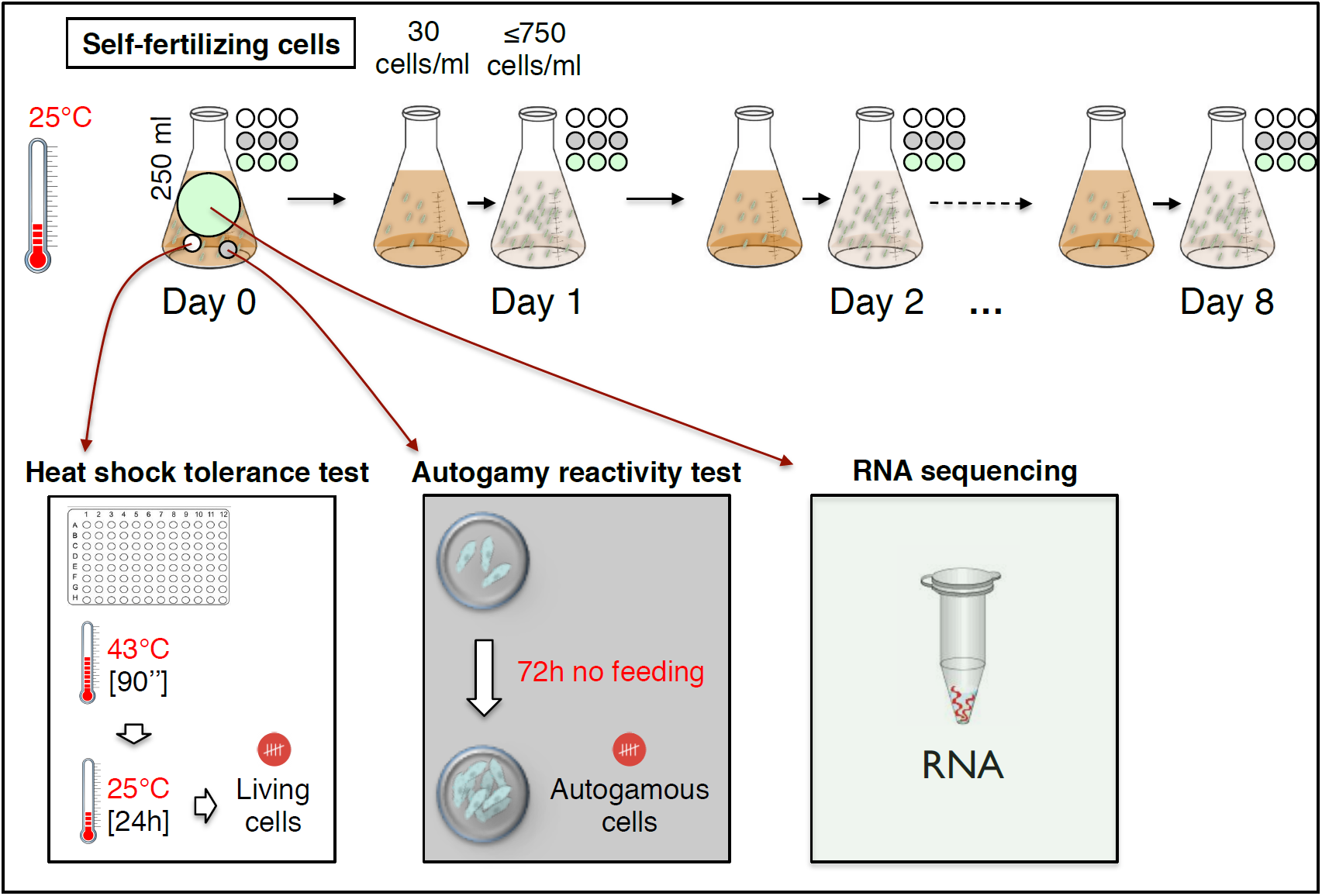
Time course experimental design.

#### Heat shock tolerance test

Mass-cultured, log-phase cells were tested daily for heat shock as they progressed through the clonal life cycle (**Figure 5**). The assay included three independent biological replicates, each consisting of 90 cells. Single cells in 1µl were transferred to a 96-well PCR plate (Art.No. 710880, Biozym Scientific GmbH) pre-loaded with 70µl/well of bacterized CM and incubated at 43°C for 1.5 min in a thermocycler (Eppendorf Mastercycler® pro S). Cells were randomly assigned to plate wells for both control (no heat shock) and treatment (heat shock) groups. Following the heat shock bout, the plates were kept at 25°C and scored for survival after 24h. Each well was labeled according to a binary classification as alive (presence of actively swimming cells) or dead (absence of actively swimming cells). To account for any asymmetry in the heat distribution within the cycler’s thermal block, we additionally estimated the binary survival classification on subsets of the plates (left and right halves). In no case, did we find differences in the pattern of heat shock tolerance.

#### Autogamy reactivity test

Mass-cultured, log-phase cells were also tested daily for autogamy reactivity (**Figure 5**). The assay included three independent biological replicates, each consisting of 56 cells on average. Briefly, 10ml aliquots were collected from triplicate mass cultures and routinely challenged with nutritional deprivation at 25°C for three days to stimulate autogamy. From each starved aliquot, cells were collected in 20µl, fixed in 0.5% (w/v) paraformaldehyde and stained with 5 µg/ml DAPI to label the nuclei. To determine the fraction of autogamous cells in each sample, at least 40 cells per replicate were screened on an epifluorescence microscope (ZEISS Axioskop 2, light source LQ-HXP 120). Cells showing intact macronuclei were scored as non-autogamous (*n*), whereas cells with extensive macronuclear fragmentation were scored as autogamous (*a*). The ratio *a*/(*a*+*n*) was taken as the fraction of autogamous cells inducible at a given age.

#### Total RNA extraction and sequencing

Total RNA was collected in triplicates for each time point (**Figure 5**). Briefly, ∼10^5^ log-phase cells from 200 ml mass culture were concentrated on hydrophilic nylon membranes (Nylon-Netzfilter, 10 µm pore size, 47 mm, Merck KGaA), rinsed in mineral water (Volvic®) and pelleted by low-speed centrifugation at 800 xg for 2 min. The pellet was immediately lysed in 1 ml of TRI Reagent® (Sigma-Aldrich, Inc.) and stored at −80°C until extraction. Total RNA isolation was performed according to the manufacturer instructions and yielded high-quality, DNA-free RNA with an average RIN of 8.3.

#### Quality control and reads preprocessing

We analyzed Illumina Pair End sequences of polyA-RNA (∼55Mb; 2×150bp)—strand-specific RNA sequencing library preparation—obtained for 27 RNA samples. Quality control of the RNA-seq data was performed using the FastQC (version 0.11.57 **[94]**). Low-quality reads were removed using default options of trim galore version 0.4.5 (https://github.com/FelixKrueger/TrimGalore). On average, more than 25 million reads per samples were produced from the sequencing.

#### Reference-guided transcriptome assembly

The RNA sequence reads were mapped with STAR version 2.5.oa (default parameters; **[95]**) onto the *Paramecium tetraurelia* genome **[96]**, with the annotation of the P51 version 2 **[75]** obtained from the *Paramecium* database (ParameciumDB) **[97]**. On average, for each sample 90% of the reads aligned to the reference genome. Transcript abundance was measured in terms of read counts using the same annotation file used for the transcriptome assembly, leveraging the featureCounts **[98]**, R Bioconductor package, default parameters. The count matrix was used as input for gene differential expression analysis.

#### Data pre-processing and gene expression visualization

A total of 31830 genes with Count Per Million (cpm) > 1 in ≥1 of the 27 samples were retained for differential gene expression analysis. We transformed the data counts using the variance stabilizing transformation function as in the Deseq2 package **[99]**. We used Principal Component Analysis (PCA) to visualize the between-sample distance based on gene expression data.

#### Differential gene expression and gene set enrichment analysis

To minimize any possible approach-specific bias, we performed pairwise differential gene expression analysis using Deseq2 **[99]** and EdgeR **[100]**. The raw count matrix was employed as input for the Deseq2 package. Concerning the EdgeR tool, the input data matrix was normalized using the TMM (trimmed mean of M values) method. After obtaining the differential gene expression tables from Deseq2 and EdgeR, we extracted the differentially expressed genes with adjusted *P*-value < 0.05 based on the Benjamini & Hochberg method **[101]** and considered only genes that were identified as differentially expressed (after correction for multiple testing) by EdgeR and Deseq2 at the same time. The functional annotation tool DAVID **[81]** and the PANTHER classification system **[102]** were used for functional enrichment analyses.

#### Extreme gene expression levels and gene set enrichment analysis

For each gene, we computed the maximum (MAX) and minimum (MIN) expression values across all the replicates and time points, from TMM based normalized gene expression data. For this analysis, we considered genes with the sum of normalized read count in all 27 samples > 30. Within a given time point, a gene was included in the MAX list if its expression value is the highest value across all time points or differ from the highest value for less than 10%. Genes that for a given time point have at least two replicates in the MAX_10%_ list were used to investigate the *most* active biological processes at a given time point with DAVID **[81]** and PANTHER **[102]**. The same logic and tools were used to create a MIN_10%_ list of genes and investigate the *least* active biological processes at a given time point.

### Cell proliferation and sexual maturation

Outside of the time-course experiment, we performed a meta-analysis of data obtained from independently propagated cell lines of three different *P. tetraurelia* strains, d12, d4-2, and 51, fed with either *Alcaligenes faecalis* (KL) — identified through sequencing of 1399 bp of the 16S ribosomal gene, this study — or *Enterobacter aerogenes* (EA) Hormaeche and Edwards (ATCC® 35028TM) (3 strains x 3 biological replicates for EA + 2 strains x ≥3 biological replicates for KL). For all lines we determined the cell proliferation rate and the age at autogamy maturity. The mean proliferation rate in the 24 hours was determined over a period of 7 to 8 days. A linear regression was fitted to the Autogamy∼Age values. Age at (auto)maturity was extrapolated from the regression equation and taken as the age (expressed in Day Post Autogamy, DPA) at which 90% of the cells were capable of undergoing autogamy when challenged with nutritional deprivation. The data were used to evaluate the relationship between cell proliferation rate and time at autogamy maturity.

### Dietary restriction and survival to stress

Outside of the time-course experiment, we also assessed the effect of diatary restriction on survival upon prolonged exposure to lethal levels of heat. Briefly, mass cultures of post-autogamous cells grown in 50 ml falkon tubes were randomly assigned to two dietary regimes, one consisting of nutrionally deprived cells and the other of cells fed with a nutrient rich diet (bacterized medium). Nutritionally deprived cells were starved at the age of six divisions post autogamy for 3 to 5 days. Inability to undergo autogamy following starvation was confirmed with DAPI staining. Fed cells were kept in exponential growth (log phase) by daily transfer in fresh, nutrient-rich medium. During the assay, both groups were exposed to 42°C in a water bath until total cell death. The experiment was performed on 12 independent biological replicates for each dietary group and cell density measured at 10-min intervals throughout the assay. Three aliquotes (technical replicates) of 50 to 100µl were taken from each mass culture and cells were counted manually under a dissecting microscope to estimate cell density. Culture density was normalized over the initial cell density recorded before the test.

## Acknowledgments

We thank Franz Goller, Florian Horn, Sarah Schaack, and Le Anh Nguyen Long for their comments on an earlier draft of this manuscript. We would also like to thank the INCIPIT Program and Marie Curie Co-fund fellowship to AST. This work was partly supported by the Deutsche Forschungsgemeinschaft (DFG, German Science Foundation) research grant to FC [CA1416/ 1-1] and carried out within the DFG Research Training Group 2220 ‘Evolutionary Processes in Adaptation and Disease’ at the University of Münster [281125614/GRK 2220].

## Author contributions

A.S.T and V.V.: *performed experiments and measurements, analyzed the data, and wrote the manuscript*. M.R.G. *helped supervise the project, provided expertise and feedback*. F.C. *conceived and designed the project, analyzed the data, supervised, wrote the manuscript, and secured funding*.

## Declaration

The authors declare that no competing interests exist.

## Availability of data and materials

All sequenced data have been deposited in the European Nucleotide Archive under accession PRJEB33070.

